# Ultrastructure expansion microscopy of axonemal dynein in islet primary cilia

**DOI:** 10.1101/2024.09.06.611752

**Authors:** Xinhang Dong, Jeong Hun Jo, Jing Hughes

## Abstract

Primary cilia are vital sensory organelles whose structures are challenging to study due to their solitary nature and intricate cytoskeleton. Current imaging modalities are limited in their ability to visualize structural details that are important for understanding primary cilia function. Ultrastructure expansion microscopy (U-ExM) is a recent superresolution imaging technique that physically expands biological specimens using a swellable hydrogel, allowing structural interrogation of small cellular components such as cilia. In this study, we apply U-ExM to mouse and human pancreatic islets to visualize the axonemal cytoskeleton and associated proteins in primary cilia. Our study reveals the expression of axonemal dynein in islet primary cilia and centrioles, with DNAI1 being a principal subunit which we validate using targeted shRNA knockdown. We conclude that U-ExM is suitable for localizing protein expression in pancreatic islet cilia which contain axonemal dynein.

## 1 INTRODUCTION

Primary cilia are microtubule-based cellular projections that control sensory and signaling functions in eukaryotic cells. In the endocrine system, primary cilia regulate pancreatic development and islet cell function, including insulin secretion which controls whole-body glucose usage (Cano et al. 2004; Hughes et al. 2020; Sanchez et al. 2022; Volta et al. 2019). As solitary sensory antennae of the cell, primary cilia have a long and thin shape that makes them challenging to capture by imaging. Yet, the molecular architecture of cilia requires characterization to understand their intricate structure-function relationship.

A number of recent studies have examined the axonemal organization of primary cilia using ultrastructural imaging in human and rodent pancreatic islets (Müller et al. 2020, 2024; Polino et al. 2023; Sviben et al. 2024; Xu et al. 2021), as well as in non-islet tissues (Kiesel et al. 2020; Sun et al. 2019). These works have updated primary cilia structure models by demonstrating intrinsic diversity of microtubule arrangements in these sensory organelles, apart from their conventional 9+0 scheme. Functionally, the classification of primary cilia as immotile structures has come under reevaluation following experimental observations of glucose-regulated primary cilia movements in islets *ex vivo* and *in vivo* (Cho et al. 2022; Li et al. 2022; Melnyk et al. 2025). Toward clarifying the nature of primary cilia motility and identifying the molecular players, studies in mouse and human pancreatic islets have demonstrated gene and protein expression of motor proteins previously thought exclusively expressed in motile cilia, principally axonemal dynein (Cho et al. 2022; Sviben et al. 2024). Given the unexpected nature and functional implications of these observations, a systematic, high-resolution examination of axonemal dynein expression in islet primary cilia is necessary for understanding the structural basis for their motility.

Here we present an imaging analysis of mouse and human islet primary cilia by ultrastructure expansion microscopy (U-ExM), focusing on the expression and distribution of major subtypes of axonemal dynein proteins. U-ExM is a chemical technique that leverages isotropic sample expansion to gain spatial resolution in protein localization, a method increasingly used in cilia and centriole research (Chong et al. 2020; Langner et al. 2024; Laporte et al. 2024; Soukup et al. 2024). Here we use the U-ExM protocol established for 2D cultured cells by the Guichard and Hamel group (Gambarotto et al. 2019; Gambarotto, Hamel, and Guichard 2021) and optimize it for 3D pancreatic islet tissue to visualize endogenous ciliary proteins by antibody labeling. Our results show that islet cells are amenable to the U-ExM procedure and that both mouse and human islet primary cilia harbor major subunits of axonemal dynein including light, intermediate, light intermediate, and heavy chains. Axonemal dynein intermediate chain 1 (DNAI1) as a core component of the dynein motor complex (Canty et al. 2021; Goodenough and Heuser 1985) is strongly detectable in islet primary cilia and centrioles in a ring-like symmetry. We validate DNAI1 staining using adenoviral-mediated gene knockdown and control staining in human airway motile cilia. Overall, our study confirms the presence of the axonemal dynein motors in islet primary cilia and centrioles and demonstrates the utility of U-ExM for resolving ciliary structural details in 3D tissue samples across species.

## 2 METERIALS AND METHODS

### Antibodies and validation

To ensure rigor and reproducibility, we used exclusively commercially available and validated primary and secondary antibodies for this study (**Table 1**). Antibodies were selected based on 1) published literature review, 2) vendor validation by affinity purification and knockout (KO)/knockdown (KD), and 3) experience in our own laboratory. Antibody performance was first assessed in unexpanded cultured MIN6 mouse beta cells prior to use in expanded samples and in mouse and human islet tissue. We assessed antibodies to 5 different axonemal dynein subunits including DNAI1, DNAI2, DNAH11, DNAL1, and DNALI1, encompassing intermediate, heavy, light, and light intermediate chains. The axonemal marker acetylated alpha tubulin (AcTUB) and centrosomal protein FGFR1OP (FOP) served as cilia and centriole references. The best-performing dynein antibody DNAI1 was additionally validated in this report using shRNA-mediated knockdown and using control motile cilia staining in human airway tissue.

**TABLE 1.**
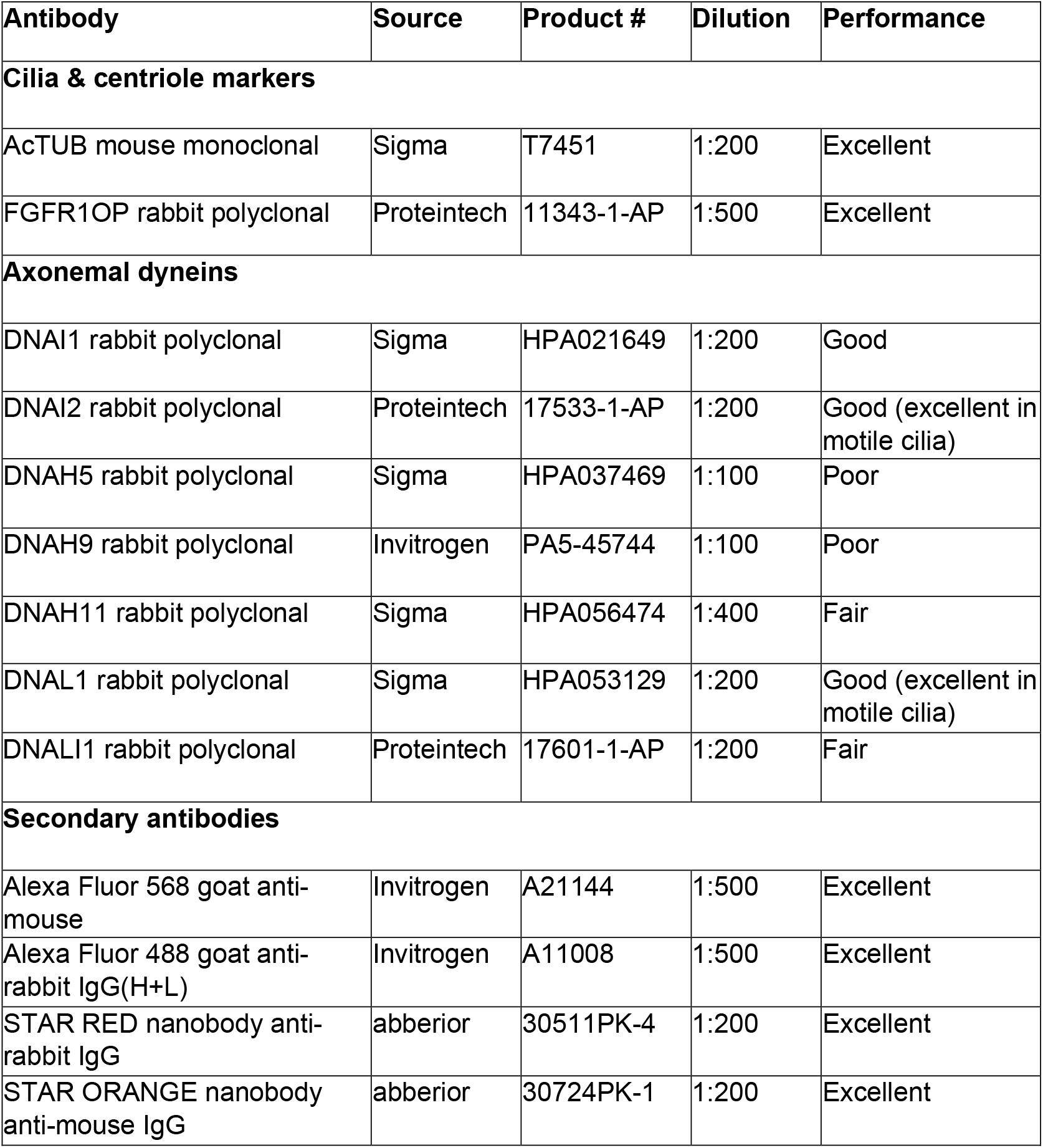
Antibody source, dilution, and performance.

| Antibody | Source | Product # | Dilution | Performance |
| --- | --- | --- | --- | --- |
| **Cilia & centriole markers** |  |  |  |  |
| AcTUB mouse monoclonal | Sigma | T7451 | 1:200 | Excellent |
| FGFR1OP rabbit polyclonal | Proteintech | 11343-1-AP | 1:500 | Excellent |
| **Axonemal dyneins** |  |  |  |  |
| DNAI1 rabbit polyclonal | Sigma | HPA021649 | 1:200 | Good |
| DNAI2 rabbit polyclonal | Proteintech | 17533-1-AP | 1:200 | Good (excellent in motile cilia) |
| DNAH5 rabbit polyclonal | Sigma | HPA037469 | 1:100 | Poor |
| DNAH9 rabbit polyclonal | Invitrogen | PA5-45744 | 1:100 | Poor |
| DNAH11 rabbit polyclonal | Sigma | HPA056474 | 1:400 | Fair |
| DNAL1 rabbit polyclonal | Sigma | HPA053129 | 1:200 | Good (excellent in motile cilia) |
| DNALI1 rabbit polyclonal | Proteintech | 17601-1-AP | 1:200 | Fair |
| **Secondary antibodies** |  |  |  |  |
| Alexa Fluor 568 goat anti-mouse | Invitrogen | A21144 | 1:500 | Excellent |
| Alexa Fluor 488 goat anti-rabbit IgG(H+L) | Invitrogen | A11008 | 1:500 | Excellent |
| STAR RED nanobody anti-rabbit IgG | abberior | 30511PK-4 | 1:200 | Excellent |
| STAR ORANGE nanobody anti-mouse IgG | abberior | 30724PK-1 | 1:200 | Excellent |

### MIN6 cells

The MIN6 cell line is a mouse pancreatic beta cell line that retains mature beta cell characteristics and expresses primary cilia (Ishihara et al. 1993; Wu et al. 2021). We used low-passage MIN6 cells, a gift from the Piston Lab at Washington University. Cells were used between P48 and P55 and maintained in DMEM media supplemented with 25 mM glucose, 15% fetal bovine serum (FBS), 1% (v/v) penicillin/streptomycin, and 4 mM glutamine.

### Animals

Wild-type mice on the C57BL/6 background were used as source of mouse islets for this study. We saw no sex differences in islet cilia morphology, number, and distribution, thus male-female data were pooled for quantitation. Animals were maintained in accordance with Institutional Animal Care and Use Committee (IACUC) regulations at the Washington University School of Medicine.

### Isolation of mouse islets

Mouse islets were isolated from healthy young adult mice by pancreatectomy followed by collagenase digestion using modified Lacy protocol (Lacy and Kostianovsky 1967; Li et al. 2009). Isolated islets were recovered overnight in islet media (RPMI-1640 supplemented with 11 mM glucose, 10% FBS, 1% (v/v) penicillin/streptomycin, and 20 mM HEPES) prior to use in experiments. Low-retention plastic and glass surfaces including tissue culture dishes and pipette tips were used to prevent islet sticking and sample loss (Stendahl, Kaufman, and Stupp 2009; Phelps et al. 2017).

### Human islets

Healthy non-diabetic human donor islets were sourced from the Integrated Islet Distribution Program (IIDP) and Prodo Laboratories Inc. Islets were inspected for morphology upon arrival and recovered overnight in islet media prior to use (RPMI-1640 supplemented with 10% FBS, 11 mM glucose, 1% (v/v) penicillin/streptomycin, 20 mM HEPES). Both male and female donor islets were included the study. Human islet use was in adherence with institutional review board (IRB) guidelines.

### Expansion microscopy

#### Protocol source

We adapted our expansion protocol from the U-ExM protocol by Paul Guichard Lab (Laporte et al. 2022), with modifications in consultation with Jennifer Wang Lab at Washington University (gelation workflow, denaturing buffer recipe) (Langner et al. 2024) and with Mu He Lab at Hong Kong University (coverslip stacks in gelation mold for 3D tissue expansion) (Chong et al. 2020).

#### Sample preparation for ExM

For 2D monolayer cells, MIN6 cells were seeded and grown to approximately 70% confluence on 12 mm glass coverslips coated with poly-D-lysine (100 μg/mL) to promote cell attachment. Cells were fixed with 4% paraformaldehyde (PFA) in PBS for 15 minutes at room temperature, permeabilized in 0.3% Triton X-100 in PBS for 10 minutes, washed in PBS, then incubated with freshly prepared acrylamide/formaldehyde (AA/FA) mix (final concentration 0.7% formaldehyde, 1% acrylamide in PBS) for 5-8 hours at 37°C.

For 3D islet tissue, intact isolated murine and human islets were rinsed in PBS and fixed in 4% PFA for 30 minutes, permeabilized in 0.3% Triton X-100 in PBS for 10 minutes at room temperature, then treated with AA/FA mix overnight at 37°C post fixation. A total of 150-200 islets were seeded on each 12 mm coverslip for expansion. Islets were over-seeded to account for attrition during sample prep.

#### Expansion workflow

A humidified gelation chamber was prepared using a 10 cm petri dish with a wet paper towel tucked around its inside perimeter. Two stacks of square coverslips were used as buttresses to support the gelation mold to ensure even thickness of the gel, which was particularly helpful for 3D tissue expansion. Glass slides were placed above and below the sample to ensure a smooth and clear surface on both sides of the gel (**Supplemental Figure 1A**). <u>Gelation</u> was performed using fresh-made gelation buffer (19% sodium acrylate, 10% acrylamide, 0.1% bis-acrylamide, 0.5% ammonium persulfate (APS), and 0.5% tetramethyl ethylenediamine (TEMED)), and cover the top slide on. Samples were incubated at 4°C on ice for 5 minutes and then at 37°C in an incubator for 45 minutes for monolayer cells and 1 hour for islet tissue. <u>Denaturation</u> was performed on solidified gels, freed from supporting slides and coverslips, and submerged in denaturation buffer (200 mM sodium dodecyl sulfate (SDS), 200 mM NaCl, 50 mM Tris-HCl pH 9) at 95°C for 1 hour. <u>Expansion</u> was performed by immersing gel samples in Milli-Q water, 30 minutes x 5 exchanges, total 150 minutes. Expanded samples were stored either in Milli-Q water in full-expanded form or cut into small pieces and stored in PBS up to 2 months. Storage in PBS shrunk the gel samples by approximately 50% in diameter which required a re-hydration and re-expansion step with 30 minutes x 2 exchanges in Milli-Q water before use.

#### Expansion factor calculation

The expansion factor is determined with experimental imaging data using centriole diameter and axial length as reference (**Supplemental Figure 1B**-**C**). The formulas for calculation are as follows:

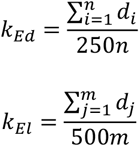

In these equations, *k*_*Ed*_ is the expansion factor of centriole diameter, and *k*_*El*_ is the expansion factor of centriole axial length. *n* and *m* represent the numbers of centrioles counted; *d*_*i*_ is the diameter of the *i*^*th*^ centriole while *d*_*j*_ is the axial length of the *j*^*th*^ centriole. The units of *d*_*i*_ and *d*_*j*_ are nanometers (nm), as is for the constant 250 which represents the average diameter of an unexpanded centriole (250 nm). The axial length of an unexpanded centriole ranges 300 to 700 nm depending on cell cycle status, thus the median length 500 nm is used as a reference for expansion factor analysis (Sahabandu et al. 2019; Uzbekov, Garanina, and Bressac 2018).

We empirically measured diameter and axial length of centrioles from images of expanded cells using both primary mouse islets (**Supplemental Figure 1**) and cultured MIN6 cells (**Supplemental Figure 2**). We used the built-in tool Full Width at Half Maximum (FWHM) in the Nikon NIS-Element software to determine centriole dimensions. The measured fluorescence intensity across the centriole produces a trapezoidal impulse wave, quantifiable as diameter and length measurements d_i_ or d_j_, which were used in above formulas for expansion factor calculation.

### Immunostaining

Post-expansion gels were punched into 8 mm circular disks using the tops of P1000 pipette tips and transferred to Eppendorf tubes for staining. Gels were first blocked in PBSBT (PBS with 3% bovine serum albumin (BSA), 0.1% Triton X-100, 0.02% sodium azide) for 30 minutes at room temperature, then incubated in primary antibodies diluted in PBSBT for 48 hours at 4°C. Samples were washed 5 times with PBSBT, 10 minutes each, then incubated with second antibodies and 4”,6-diamidino-2-phenylindole (DAPI, 0.5 μg/mL) diluted in PBSBT overnight at 4°C. Stained samples were washed in Milli-Q water for 15 minutes x 5 times, then stored in Milli-Q water protected from light. Stained samples were imaged within two weeks of preparation.

### Confocal and super-resolution microscopy

Immunofluorescence imaging was performed on a Zeiss LSM 880 confocal microscope and a Nikon AXR super-resolution imaging system with Nikon spatial array confocal (NSPARC). For sample stability, stained and expanded gel samples were placed in a 35 mm glass-bottom dish coated with poly-D-lysine to minimize sample movement and the sample surrounded with a wet paper towel to prevent gel drying and shrinking. Imaging was done using 20x, 60x, and 100x oil objectives to obtain a range of magnifications.

### Transmission Electron Microscopy (TEM)

Primary cilia and basal bodies from mouse and human islets were imaged at ultrastructural resolution using TEM. Intact islets were incubated overnight in RPMI islet media, fixed in a modified Karnovsky”s solution containing 3% glutaraldehyde and 1% paraformaldehyde in 0.1M sodium cacodylate buffer (SCB) at 4°C overnight, then fixed in 2% osmium tetroxide in 0.1M SCB for 1 hour, en bloc stained with 3% aqueous uranyl acetate for 30 min, dehydrated in graded ethanol, and embedded in PolyBed 812 (Polysciences, 08792-1). Islet pellet sections of 90-nm thickness were cut, stained with Venable”s lead citrate, and examined with a JEOL 1200 EX. Digital images were acquired using the AMT NanoSprint12 (Advanced Microscopy Technology) CMOS, 12-megapixel TEM camera.

### Adenoviral knockdown

Dynein axonemal intermediate chain 1 (*Dnai1*) gene knockdown was performed in mouse islets using adenoviral particles (pAV[shRNA]-EGFP-U6>mDnai1[shRNA#1], Vector ID: VB240719-1668mrt, VectorBuilder). Control knockdown was scramble shRNA control adenovirus (Vector ID: VB010000-0020ytx, VectorBuilder). A minimum of 50 islets per mouse (n=3 mice) were used per knockdown condition. Intact islets were transduced without dissociation with either control or *Dnai1* shRNA adenoviral particles at 1 × 10^8 infectious unit (IFU)/mL for 24 hours at 37°C. Post-shRNA transduction, islets were recovered in a 35 mm petri dish for 96 hours with media changes every other day before using for immunostaining and mRNA quantitation by qPCR.

### Quantitative polymerase chain reaction (qPCR)

shRNA knockdown efficiency was validated by qPCR. Total bulk islet mRNA was isolated from transduced mouse islets by lysing islets in 350 μL of lysis buffer RA1 (Macherey-Nagel, 740961) with β-mercaptoethanol, followed by RNA purification using the Nucleospin RNA kit (Macherey-Nagel, 740955.50). Complementary DNA (cDNA) was synthesized with High-Capacity cDNA reverse transcription kit (Thermo Fisher Scientific, 4368814). Quantitative PCR was performed in triplicate on a QuantStudio 3 Real-Time PCR Instrument (Thermo Fisher Scientific, A28131) using 2x Power SYBR Green PCR Master Mix (Thermo Fisher Scientific, 4367659). Gene expression levels were quantified using 2^−ΔΔCt^ and normalized by the housekeeping gene *Gapdh*. The following primer sequences were used: mouse *Dnai1*: F: 5”-ACCTCCAAGTCTGGCAAGCACA-3”, R: 5”-ACTATCCTGCCATCGGACGACA-3”; mouse *Gapdh*: F: 5”-AGGTTGTCTCCTGCGACTTCA-3”, R: 5”-CAGGAAATGAGCTTGACAAAGTTG-3”.

## 3 RESULTS

### 3.1 Ultrastructural imaging of islet primary cilia revealing axonemal dynein

We assessed the morphology of primary cilia and centrioles by TEM and by confocal microscopy in unexpanded and expanded pancreatic islet tissues. The primary cilium contains a microtubule-based, electron-dense axonemal cytoskeleton and an overlying ciliary membrane continuous with the plasma membrane (**Figure 1A-B**). A typical pancreatic islet primary cilium measured 3-5 μm in length and 200-300 nm in diameter, with a shallow ciliary pocket visible on cross-section by TEM and a 9-fold radial symmetry of microtubule filaments at the base (**Figure 1B**). Confocal imaging of expanded islet cilia and centriole demonstrated expected length and diameter increases corresponding to 4x isotropic expansion, and similar cytoarchitectural arrangements as seen on TEM including axonemal microtubules exhibiting a distinct radial symmetry as marked by acetylated alpha tubulin staining (AcTUB, red, **Figure 1C**), suggesting that islet cilia structures are preserved among imaging modalities.

**FIGURE 1.**
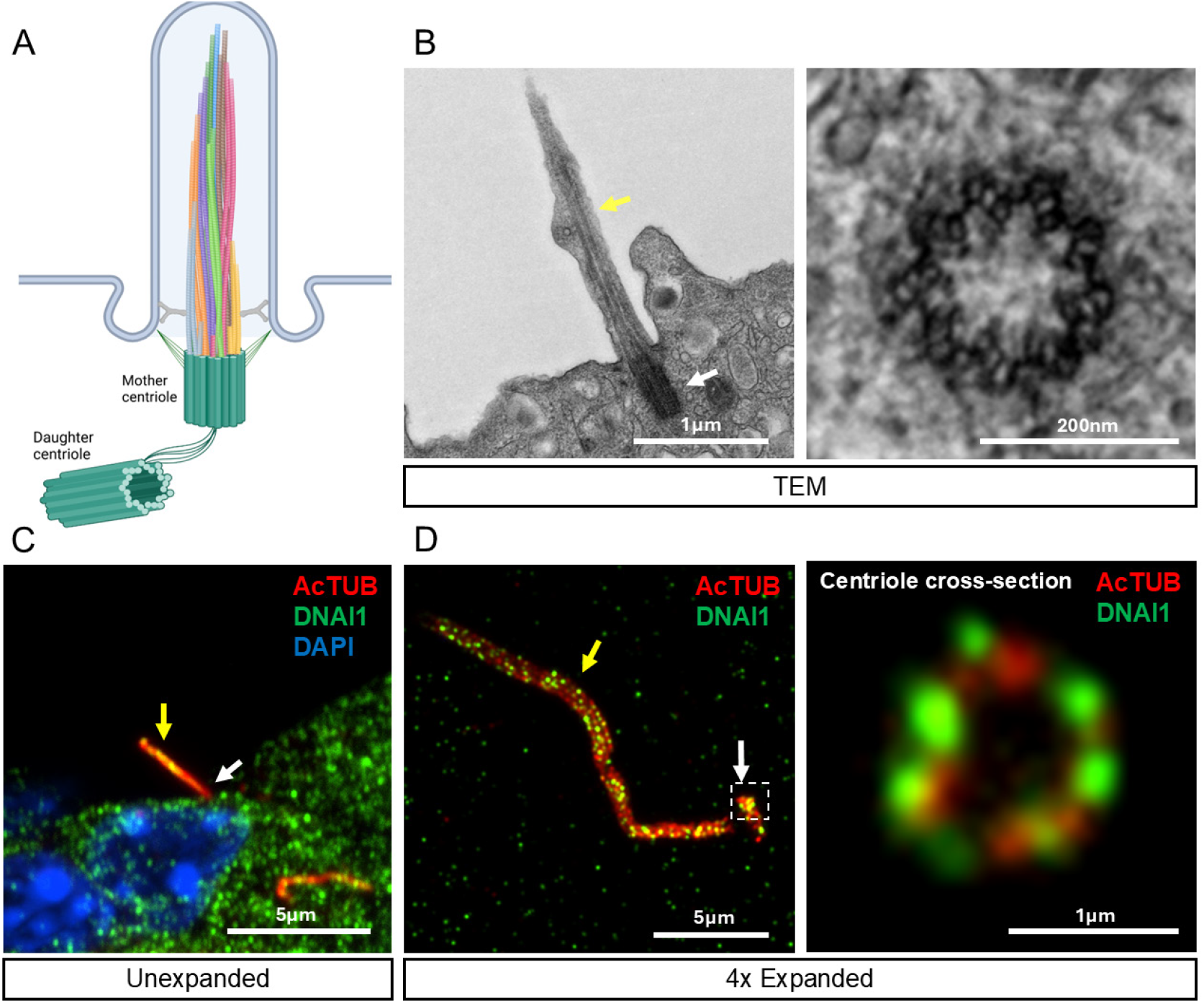
Ultrastructural imaging of pancreatic islet primary cilia. (**A**) Schematic structure of the primary cilia axoneme, depicting a rotating bundle of microtubules that evolves from base to tip. (**B**) Transmission electron microscopy (TEM) showing longitudinal view (left) of the axoneme and basal body of a mouse islet primary cilium, and cross-section view (right) of a human islet basal body or centriole showing 9+0 symmetry of microtubule triplets. (**C**) Immunostaining of axonemal dynein intermediate chain 1 (DNAI1, green) in native unexpanded mouse islets showing localization to primary cilia (AcTUB, red) and cytosol, 2 μm z-projection. (**D**) NSPARC confocal images of mouse islet post-expansion showing DNAI1 localization to the ciliary axoneme and centriole; inset showing orthogonal cross-section view of the tubulin ring (red) with DNAI1 (green) distributed radially in discrete foci on the ring periphery. Cellular dimensions are increased post-expansion as referenced by scale bars. Yellow arrows, axoneme; white arrows, basal body.

We observed endogenous expression of axonemal dynein intermediate chain 1 (DNAI1) in islet primary cilia and centrioles by confocal imaging in native unexpanded mouse islet samples (DNAI1, green, **Figure 1C**). Closer evaluation of DNAI1 expression by U-ExM in expanded mouse islets showed that the motor protein is present throughout the length of the primary cilium and centrioles, exhibiting a ring-like distribution of dynein around the tubulin filaments (**Figure 1D**). Expansion microscopy reduced the background cytoplasmic signal by DNAI1 antibody and allowed clear visualization of axonemal dynein in association with tubulin in islet primary cilia (**Figure 1D**).

### 3.2 Axonemal dynein expression is conserved between primary and motile cilia

As functional axonemal dynein complexes require four classes of subunits including light, intermediate, light intermediate, and heavy chains, we investigated the expression of all 4 types of axonemal dynein in mouse islet primary cilia, using human airway motile cilia as positive control. Among the antibodies we tested, DNAI1, I2, L1, LI1, and H11 consistently produced ciliary staining in mouse islets, in addition to punctate background cytoplasmic staining. Ciliary signals of axonemal dynein are best seen in projecting cilia at the islet periphery, away from the background signal in the cell body (**Figure 2A**). In the motile cilia of cultured primary human airway cells, axonemal dynein staining was strongly ciliary for all antibodies tested, with little to no background signal in the cell body, suggesting either tissue-dependent differences in subcellular dynein distribution or superior antibody performance in human airway cells (**Figure 2B**). Two additional heavy chain markers, DNAH5 and H9, were each detected in mouse islet primary cilia (DNAH5) and human airway motile cilia (DNAH9) but not reliably in both. As additional optimizations may be required for these two antibodies, we elected to exclude them from this report.

**FIGURE 2.**
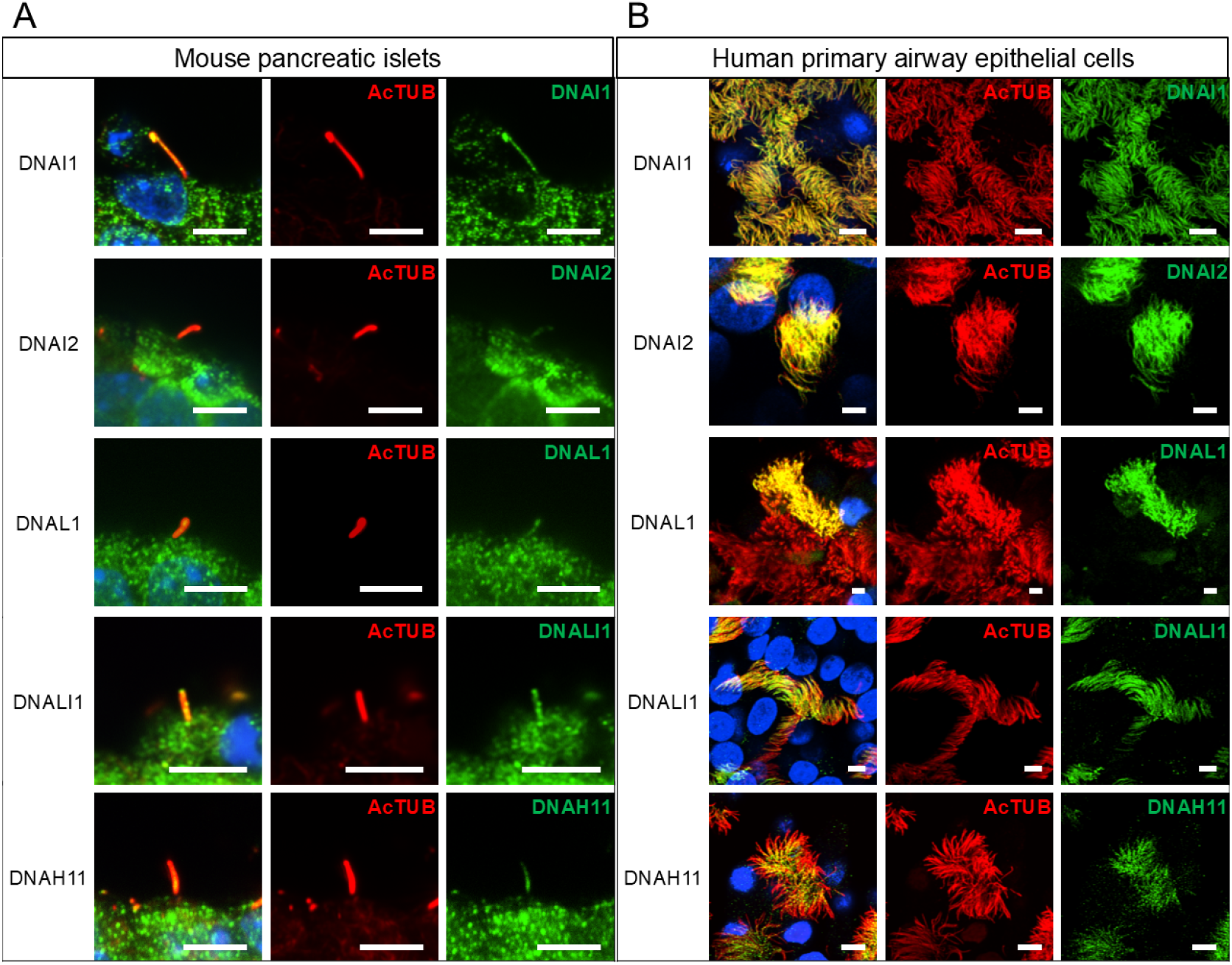
Axonemal dynein expression in primary and motile cilia. Representative confocal images of axonemal dynein DNAI1, DNAI2, DNAL1, DNALI1, and DNAH11 expression in (**A**) primary cilia (mouse islets) and (**B**) motile cilia (human airway cells). Dynein subunits are shown in the green channel, cilia marker acetylated tubulin (AcTUB) in red, and merged views with nuclei (DAPI) in blue; scale 5 μm; images are 2 μm-thickness z-projections to capture entire ciliary axonemes. While most dynein subunits are detected continuously along both the primary and motile cilia axoneme, DNAH11 is more proximally concentrated in both islet primary cilia and airway motile cilia (bottom panels).

### 3.3 Human islet primary cilia contain axonemal dyneins with known roles in PCD

Two axonemal dynein proteins were further evaluated in human islet cells, given their prominent roles in genetic human motile ciliopathies. DNAI1 as a core component of the outer dynein arm (ODA) is a common mutational cause of human primary ciliary dyskinesia (PCD) (Zariwala et al. 2006). We identified DNAI1 protein expression in human islet primary cilia in situ, overlapping with the axonemal marker acetylated alpha tubulin (**Figure 3A**). The ciliary staining of axonemal dynein in human islets stands apart from the punctate cytoplasmic background signal and is detectable on the green channel alone. Ciliary DNAI1 signal spans the axonemal length and is concentrated in the proximal and base regions, as well as in centrioles that are frequently found in pairs near the nucleus (**Figure 3B**). Centriolar DNAI1 is distributed throughout the length of the barrel in semi-regular repeats, co-localizing with acetylated alpha tubulin and appearing on the outer edge of the tubulin ring (**Figure 3C**).

**FIGURE 3.**
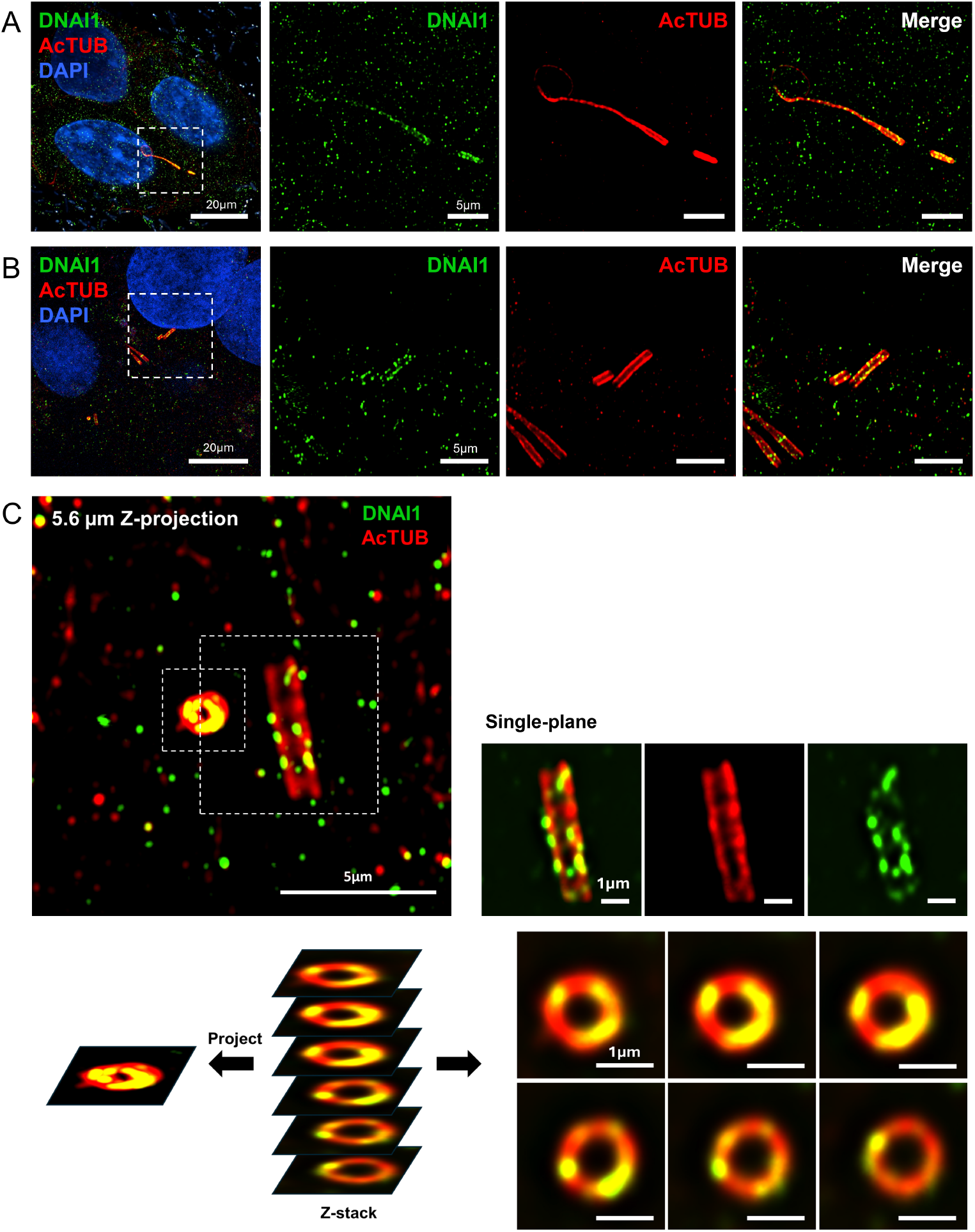
Axonemal and centriolar expression of DNAI1 in human islet cilia. Confocal z-projections (2 μm) of human islets showing axonemal dynein DNAI1 expression in (**A**) ciliary axoneme and (**B**) centriole pairs. (**C**) 5.6 μm z-projection image of human islet centrioles showing punctate staining of axonemal dynein (DNAI1, green) with the centriolar barrel marked by acetylated alpha tubulin (AcTUB, red). Scale 1-20 μm as indicated.

DNAH11 is another ciliary outer dynein arm protein that is a microtubule-dependent motor ATPase, whose mutations are found in patients with PCD and *situs inversus* (Knowles et al. 2012). We mapped its expression to the proximal end of the human islet primary cilium, most pronounced in the basal body (**Figure 4A**). In centrioles, DNAH11 is polarized toward one-half of the length of the barrel and at the distal centriolar cap (**Figure 4B-C**). These distribution patterns are consistently seen across islet cells from multiple donors, suggesting functional partitioning of this protein in human islet cilia and centriolar compartments. Both DNAI1 and DNAH11 have not previously been classified as primary cilia proteins in any cell type.

**FIGURE 4.**
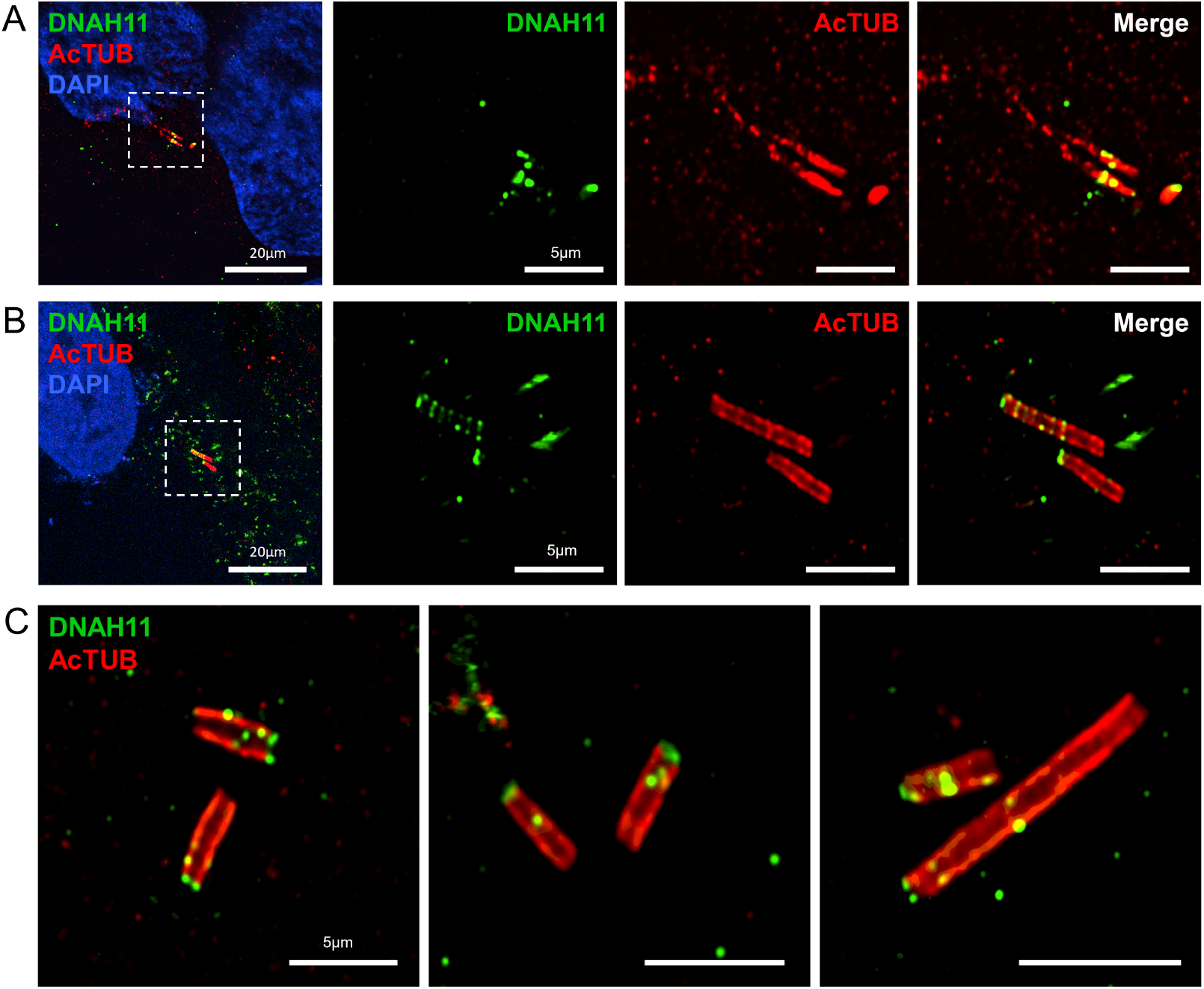
Axonemal and centriolar expression of DNAH11 in human islet cilia. Confocal z-projections (2 μm) of human islet cells post-expansion showing axonemal dynein heavy chain DNAH11 expression in the ciliary axoneme (**A**) and centrioles (**B**) in healthy human islet cells. (**C**) High-magnification images of human islet centrioles showing polarized expression of DNAH11 (green) at the centriole cap. Acetylated alpha tubulin, AcTUB (red), cilia and centriole marker. Scales 5-20 μm as indicated.

### 3.4 U-ExM visualization of DNAI1 and DNAH11 in mouse islets

To examine conservation across species and to visualize dynein expression in mouse primary cilia in higher resolution, we performed U-ExM on mouse islets and labeled for endogenous dynein subunits. Both DNAI1 and DNAH11 are detected in mouse islets post-expansion, with stronger staining in cilia than in the cell body (**Figure 5**). Axonemal dynein staining spans the length of mouse islet primary cilia, which tend to be longer than human islet cilia, and can be seen concentrated in the axonemal tip and base.

**FIGURE 5.**
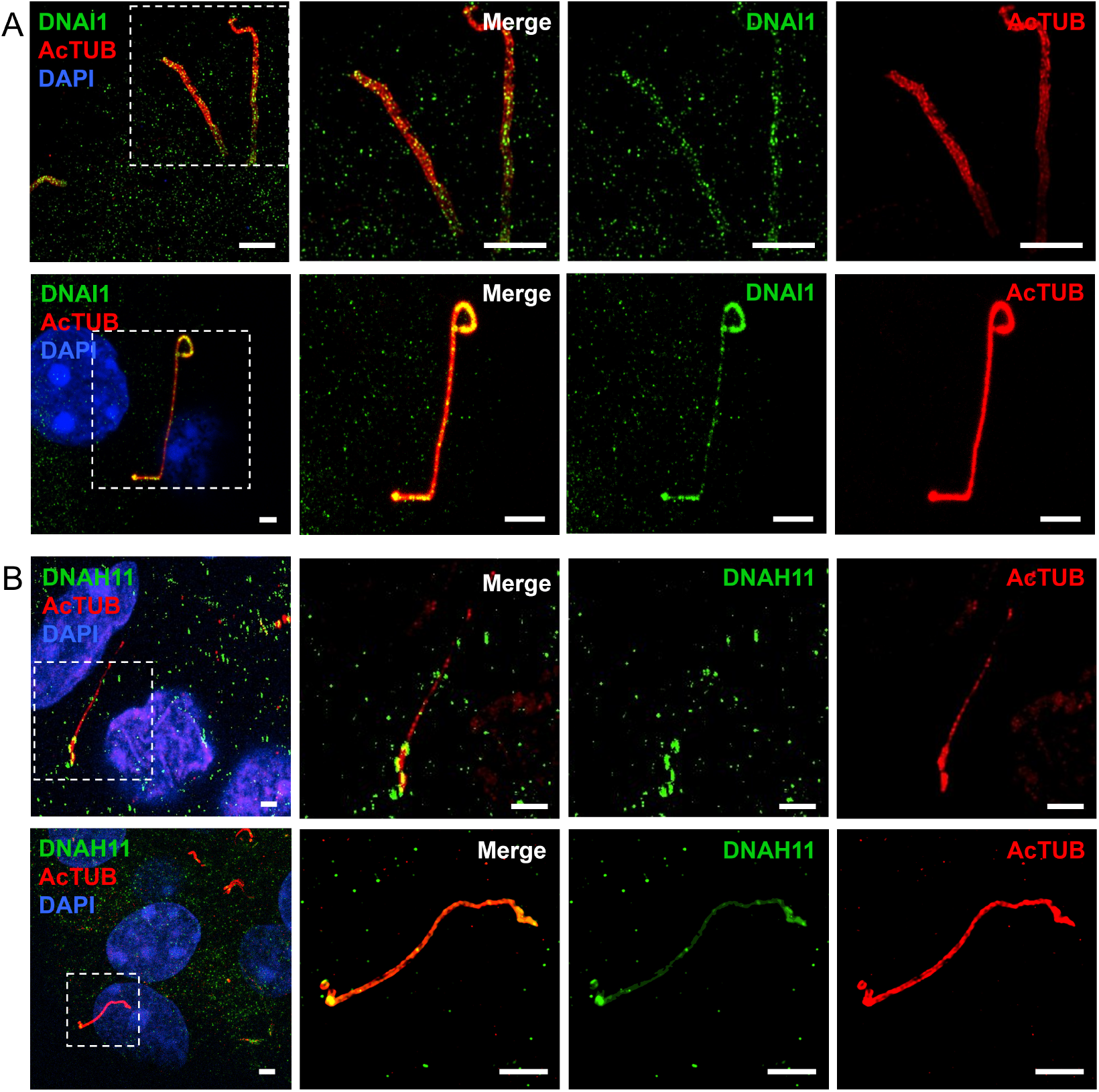
Super-resolution imaging of axonemal dynein in expanded mouse islets. Mouse islets post-expansion labeled with axonemal dynein (**A**) intermediate chain DNAI1 and (**B**) heavy chain DNAH11, two representative regions each showing z-projections of 1-5 μm thickness. Insets are displayed in merged and single-channel views. Mouse islet cilia are longer and more polymorphic than human islet cilia and tend to form loops and tendrils, seen here in both axonemal dynein (green) and acetylated tubulin (red) channels; nuclei (DAPI, blue); scale 5 μm.

These distribution patterns are observed through repeated experiments using both male and female mice and are felt representative of normal mouse islets.

### 3.5 DNAI1 staining is validated by shRNA

To ensure specificity of axonemal dynein antibody labeling, we performed adenoviral knockdown of intermediate chain gene *Dnai1* in mouse islets and repeated the DNAI1 staining. Adenovirus infection of intact non-dispersed islets achieved a 45-47% transduction efficiency as assessed by EGFP marker and 26% reduction in bulk-islet *Dnai1* mRNA as assessed by qPCR. Within EGFP^+^ transduced cells, *Dnai1* shRNA led to 76% reduction in DNAI1 protein expression compared to control shRNA in the cilium (p<0.0001), and 35% reduction in the cell body (p=0.2299, ns), suggesting ciliary selectivity of DNAI1 expression and of its shRNA knockdown (**Figure 6**). The ROI segmentation and analysis workflow for ciliary and cytosolic DNAI1 protein quantitation is presented in **Supplemental Figure 4**. Of note, because the shRNA adenovirus carried EGFP, we paired a far-red secondary antibody (abberior STAR-RED) for DNAI1 staining in knockdown experiments, instead of the green Alexa Fluor 488 secondary antibody used to stain DNAI1 in native non-transduced islets (**Figures 1-5**). Both secondary antibodies demonstrated comparable DNAI1 staining intensity and subcellular distribution, attesting to the validity of our antibody reagents.

**FIGURE 6.**
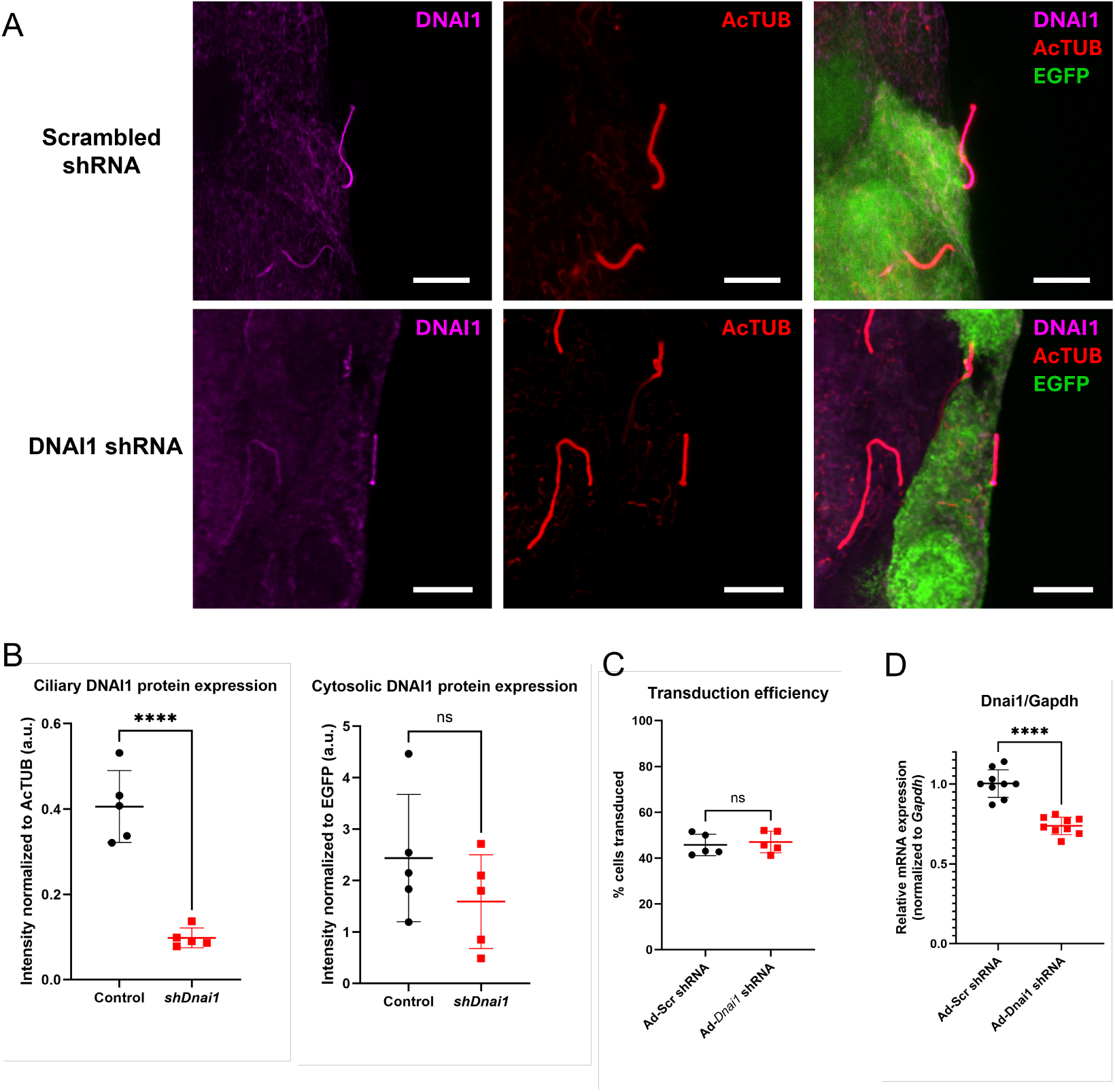
DNAI1 staining validation by targeted gene knockdown. (**A**) Mouse islets transduced with control scrambled and *Dnai1* adenoviral shRNA and stained for DNAI1 (magenta) and AcTUB (red); transduction marker EGFP (green); scale 5 μm. (**B**) Targeted knockdown reduces DNAI1 staining in cilia (left, 76%) and not in cytosolic regions (right, 35%); n = 3 averaged ROIs per islet, 5 islets per experiment, data representative of 3 experiments. (**C**) Transduction efficiency quantitation showing comparable % EGFP^+^ cells in control vs *Dnai1* shRNA transduced islets, 45% vs 47% respectively, p=06793, *ns*. (**D**) *Dnai1* knockdown quantitation by bulk-islet qPCR showing 26% reduction by *Dnai1* shRNA, triplicate islet samples measured in triplicate qPCR reactions (n = 9 per knockdown condition). One-way ANOVA, ****p < 0.0001, ns, not significant.

## 4 DISCUSSION

In summary, our study reveals localization of axonemal dynein to the primary cilium and centriole of human and mouse pancreatic islet cells. Using a combination of high-resolution confocal and expansion microscopy, we identified five of the main molecular components of axonemal dynein, one principal subunit which we validate using gene knockdown. We propose that these dynein proteins are present programmatically to enable ciliary motile function in these cells.

The demonstration of motile protein expression in human islet primary cilia corroborates prior results (Cho et al. 2022; Sviben et al. 2024). The findings of primary cilia motility and motile protein expression are unexpected and require critical data support. While mRNA expression of motile cilia genes such as DNAI1 may be low in pancreas in datasets such as Genotype-Tissue expression (GTex) Portal and Human Protein Atlas, a recent human islet transcriptome study demonstrated perturbations in motile cilia genes including axonemal dynein light, intermediate, heavy chains, dynein assembly factors, as well as the motile cilia transcription factor RFX3 to be associated with human type 2 diabetes (Walker et al. 2023). Thus, motility may be an overlooked aspect of primary cilia function in pancreatic biology, and more needs to be done to clarify the biomechanical basis and function of their motility. Whether active motility and axonemal dynein expression are a feature of primary cilia in other cell types also awaits investigation.

We additionally demonstrate centriolar expression of axonemal dynein intermediate and heavy chains DNAI1 and DNAH11 in human islet cells. Axonemal dynein proteins have not been previously reported to localize to the triplet microtubule structure of centrioles, even for canonical motile cilia. Yet, centrioles are dynamic structures within cells whose motility and migration are vitally important to cytokinesis and have also been observed in non-cell cycle settings, which mechanistically is thought to depend on plus-end directed microtubule transport, kinesin, and cytoplasmic dynein (Hannaford and Rusan 2024; Jonsdottir et al. 2010; Koonce et al. 1999; Piel et al. 2000). With increasing recognition of centrioles and centrosomes as cell signaling compartments, it would be important to understand not only how centrioles move but also how their motility contributes to signaling function. We ourselves have observed rapid and irregular centriole translocations within live pancreatic islet cells by SiR-tubulin labeling, as well as supernumerary centrioles in human islet cells (unpublished data), the biological significances of which await further study.

The number of dynein proteins examined in this study likely represents a small partial list. Axonemal dynein is a complex biological motor encompassing dozens known subunits, most of which lack reliable staining reagents. A worthwhile future direction, therefore, would be unbiased proteomic demonstration of axonemal dynein subunits in purified primary cilia, from islet cells or otherwise, which would strengthen the notion that primary cilia harbor motile machinery. Other imaging approaches including iterative U-ExM, cryo-EM mapping of individual axonemal components, and tagging these proteins *in situ* to observe their localization and trafficking patterns would provide additional information on their function in primary cilia. A detailed understanding of the distinct distributions of axonemal dynein subunits within primary cilia and centriole compartments may reveal their unique roles in the assembly and function of the motile machinery in these cellular structures.

## 5 CONCLUSION

We apply conventional confocal and expansion microscopy to study axonemal dynein expression in mouse and human pancreatic islets. The detection of motor dynein subunits in pancreatic islets across species suggests a potential conserved role for cilia motility in normal islet tissue function. Future research might explore the biomechanical basis and functional purpose of dynein-dependent cilia motility in pancreatic islets, as well as primary cilia motility in other cellular contexts.

## ACKNOWLEDGEMENT

We thank Dr. Steven Brody for providing human airway cells for motile cilia staining. We thank Drs. Jennifer Wang and Mu He for U-ExM protocol sharing and technical advice. Microscopy was performed at the Washington University Center for Cellular Imaging (WUCCI) supported by Washington University School of Medicine, The Children”s Discovery Institute of Washington University and St. Louis Children”s Hospital (CDI-CORE-2015-505 and CDI-CORE-2019-813) and the Foundation for Barnes-Jewish Hospital (3770 and 4642). Graphical illustrations created with BioRender. Funding for this study provided by NIH grants R01DK138974 and R01DK140365 to JWH.

