## Supplemental Figures 1-3 for "Ultrastructure expansion microscopy of axonemal dynein in islet primary cilia"

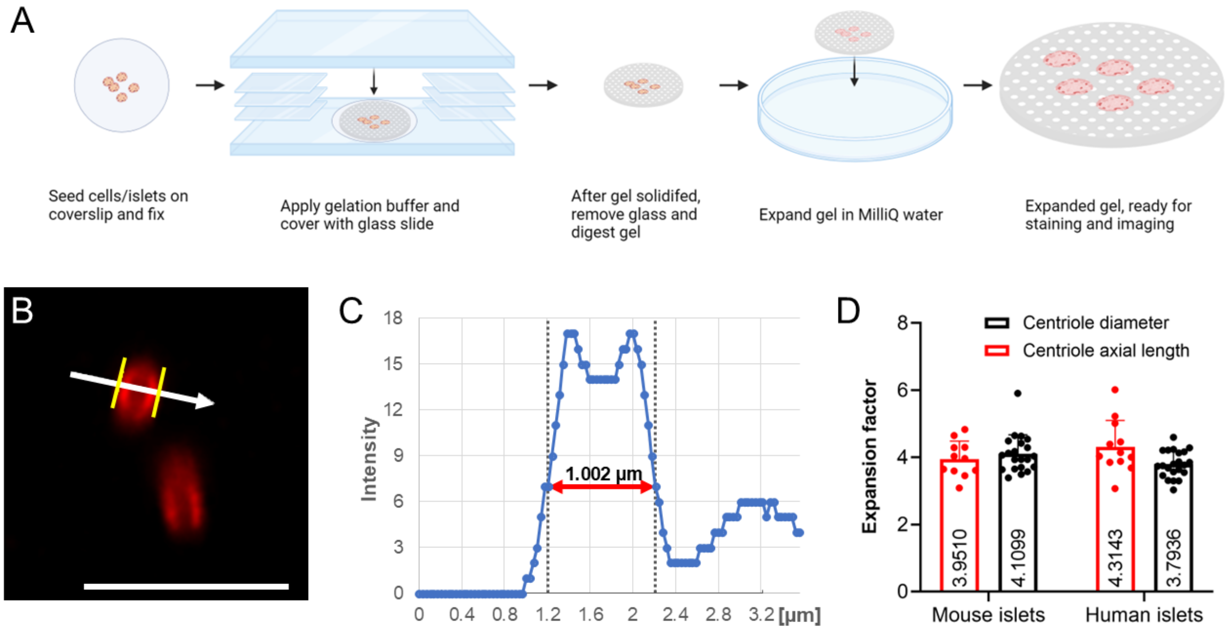


**SUPPLEMENTAL FIGURE 1. Workflow of ultrastructural expansion microscopy (U-ExM) of pancreatic islets.** (**A**) Schematic showing the seeding, gelation, expansion, and staining process. Antibody labeling is performed post-expansion, as this in our hands produced superior labeling than pre-expansion labeling. (**B**) Centriole diameter measurement in Nikon NIS-Elements software for expansion factor calculation; shown are expanded mouse islet centrioles labeled with acetylated alpha tubulin (AcTUB, red), scale 5 µm. (**C**) Full width at half maximum (FWHM) measurements of expanded mouse islet centrioles, measuring ~1 µm which corresponds to 4x expanded native centriole diameter of ~250 nm. (**D**) Expansion factors calculated from mouse and human islet centrioles. n = 21-22 for centriole diameter measurements and n = 11-12 for centriole axial length measurements. p values: mouse diameter/axial = 0.8687; human diameter/axial = 0.0539; diameter mouse/human = 0.2541; axial mouse/human = 0.4058, all non-significant, attesting to the reproducibility of the expansion procedure and analysis.

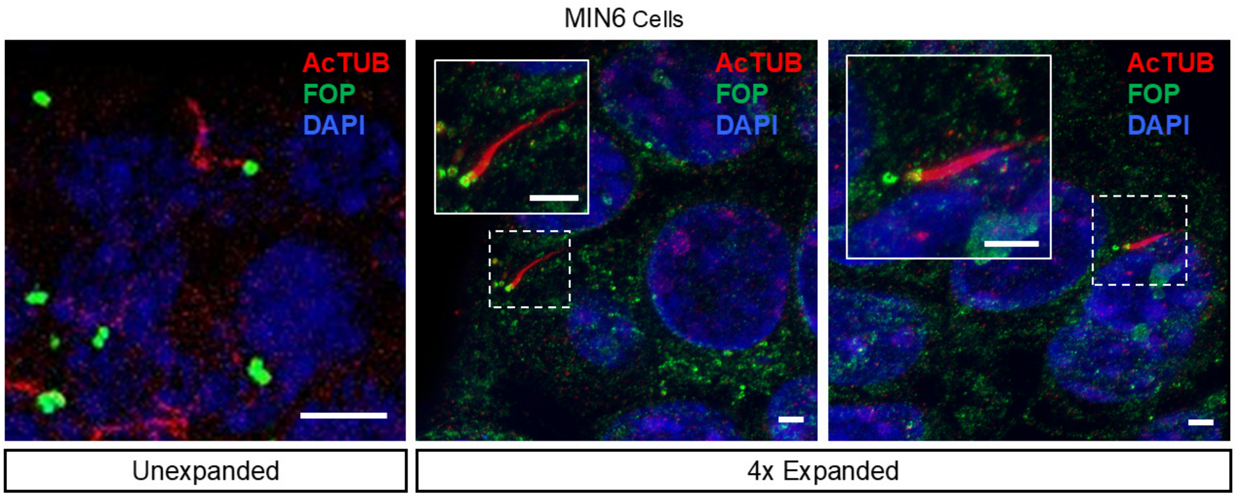


**SUPPLEMENTAL FIGURE 2. Cilia and centriole imaging in MIN6 beta cells by U-ExM.** Native unexpanded and 4x expanded MIN6 beta cells imaged on a Zeiss LSM880 confocal microscope showing preserved ciliary and cyto-morphology after expansion. Cells are co-labeled with axoneme marker acetylated tubulin (AcTUB, red), centriole marker FGFR1 oncogene partner (FOP, green), and nuclei (DAPI, blue). Solid-border insets are enlarged from dashed-border regions to show detail, scale 5 μm.

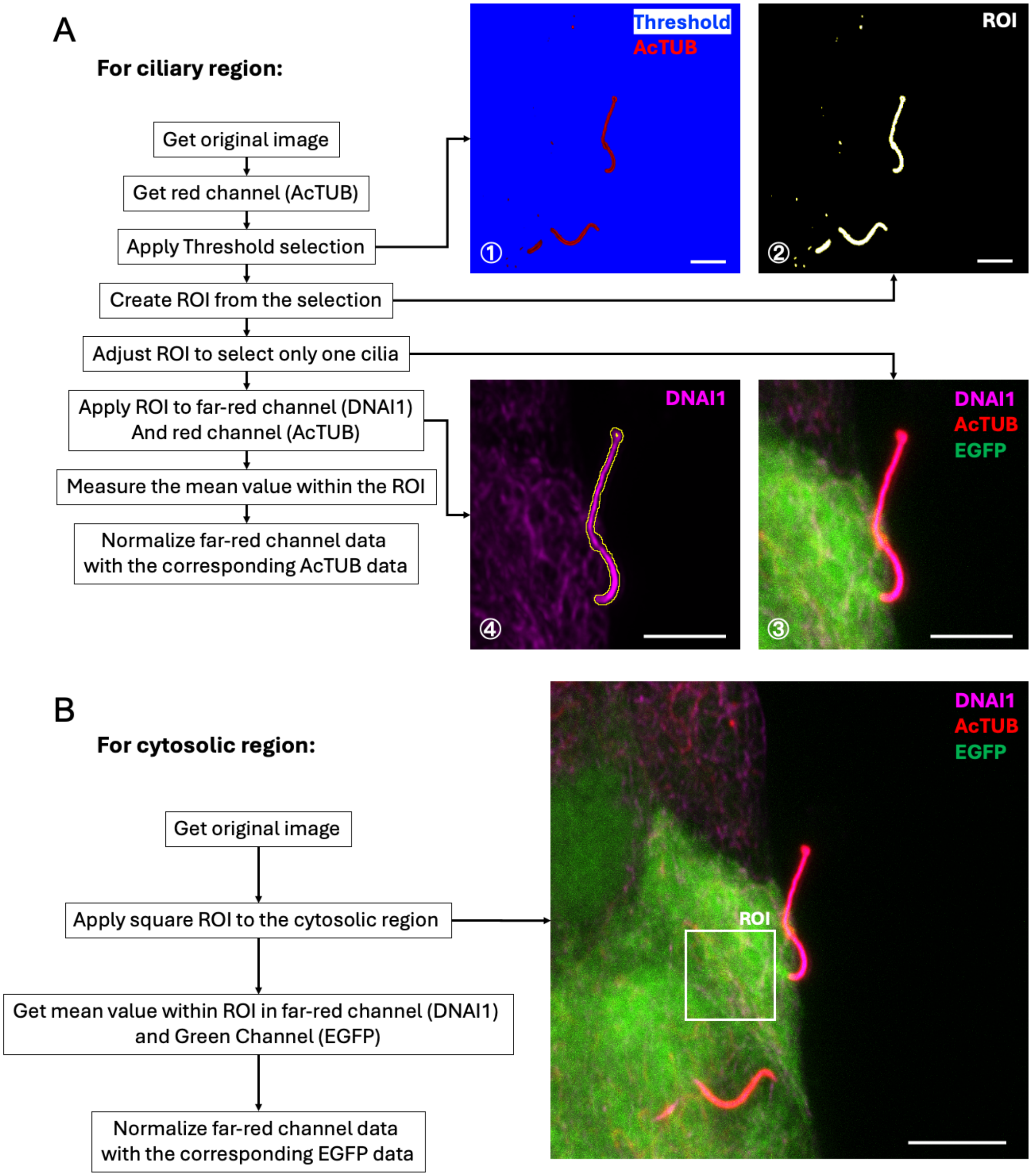


**SUPPLEMENTAL FIGURE 3. Region of interest (ROI) selection for ciliary vs cytosolic shRNA staining quantitation.**
